# *Talaromyces marneffei* promotes M2 polarization of human macrophages by downregulating SOCS3 expression and activating TLR9 pathway

**DOI:** 10.1101/2021.02.17.431726

**Authors:** Wudi Wei, Chuanyi Ning, Jiegang Huang, Gang Wang, Jingzhen Lai, Jing Han, Ning Zang, Bingyu Liang, Yanyan Liao, Thuy Le, Junjun Jiang, Li Ye, Hao Liang

## Abstract

Little is known about how *Talaromyces marneffei*, a thermally dimorphic fungus that causes substantial morbidity and mortality in Southeast Asia, evades the human immune system. Polarization of macrophages into fungal-inhibiting M1 and fungal-promoting M2 types has been shown to play an important role in the innate immune response against fungal pathogens. This mechanism has not been defined for *T. marneffei*. Here, we demonstrated that *T. marneffei* promotes its survival in human macrophages by inducing them towards M2 polarization. Our investigations of the mechanism revealed that *T. marneffei* infection led to SOCS3 protein degradation by inducing tyrosine phosphorylation, thereby relieving the inhibitory effect of SOCS3 on p-STAT6, a key factor for M2 polarization. Our SOCS3-overexpression experiments showed that SOCS3 is a positive regulator of M1 polarization and plays an important role in limiting M2 polarization. Furthermore, we found that inhibition of TLR9 pathway partially blocked *T. marneffei*-induced M2 polarization and significantly enhanced the killing activity of macrophages against *T. marneffei*. Collectively, these results reveal a novel mechanism by which *T. marneffei* evades the immune response of human macrophages.

## Introduction

*Talaromyces marneffei* (formerly *Penicillium marneffei)* is a thermally dimorphic fungus endemic in southeast Asia, southern China, and northeastern India and causes an invasive fungal infection in immunocompromised individuals living in or traveling to these endemic regions ^1,2^. Human infection is presumed to occur in the lung via inhalation of aerosolized conidia from the environment. The fungus is able to evade the host immune response to establish a latent infection which, in the setting of immunosuppression, can reactivate within the reticuloendothelial system to cause a life-threatening multi-organ disseminated infection ^2^. Advanced HIV disease (CD4 cell count <100 cells/mm^3^) is a major risk factor for talaromycosis. In the highly endemic regions of northern Thailand, Vietnam, and southern China, *talaromycosis* is diagnosed in up to 16% of HIV hospital admissions, and is a leading cause of HIV-associated death ^3,4^. Infections are increasingly diagnosed in non-HIV-infected people who have a primary immunodeficiency condition (such as idiopathic CD4 lymphopenia, anti-interferon-gamma autoantibodies, mutations in the CYBB, CD40L, or STAT pathways) and in those who have a secondary immunodeficiency condition (corticosteroids or immunosuppressive therapy, malignancies, and solid or bone marrow transplantation) ^5^. The mortality despite antifungal therapy is up to 30% in HIV-infected individuals and up to 50% non-HIV-infected individuals ^6–8^.

Despite high morbidity and mortality and a growing number of susceptible individuals, our current knowledge of the pathogenicity of *T. marneffei* causing human disease is limited. The strategy in which *T. marneffei* evades the innate immune response in the lung is unknown. Specialized phagocytes of the innate immune system provide the first line of defense against fungal infection ^9^. The activation of macrophages is critical for their anti-fungal function. Macrophages are activated into two functionally distinct M1 and M2 phenotypes ^10^. M1 macrophages are associated with an immune response to intracellular pathogens and are involved in pro-inflammatory responses governed by Toll-like receptors (TLRs) and Th1 interferon gamma signaling. M2 macrophages are associated with an immune response to helminth infections and asthma and allergies and are involved in anti-inflammatory responses and tissue repair governed by Th2 signaling ^10^. *T. marneffei* infection has been shown in a mouse alveolar macrophage model to induce phenotype M2-associateded factor, IL-10 ^11^. Another study using human macrophage model has shown that *T. marneffei* infection up-regulates the level of IL-10, a M2 macrophage-associated cytokine with the main function is to inhibit the antifungal activity of macrophages ^12^. A recent study using zebrafish model showed that macrophages protect *T. marneffei* conidia from neutrophil antifungal activity; however, the specific mechanism is unknown ^13^. Since the strategy for immune evasion by promoting macrophage M2 polarization has been proposed for other intracellular pathogens including *Candida albicans* and *Staphylococcus aureu* ^14–16^, we hypothesize that *T. marneffei* may evade macrophage antimicrobial functions similarly by inducing M2 polarization.

Previous studies have identified several proteins and pathways that regulate macrophage polarization, including suppressor of cytokine signaling (SOCS) family and JAK-STAT pathway ^10,17,18^. Among SOCS family, SOCS3 has been shown to regulate M1 polarization in macrophage ^19^, but this regulation may be species-specific ^18^. For example, SOCS3 functions as a negative regulator of the M1 polarization in a mouse macrophage model ^20–22^, but functions as a positive regulator of the M1 phenotype in a rat bone marrow-derived macrophages (BMM) and a human THP-1 macrophage model ^19,23^. The JAK-STAT6 pathway is involved in the M2 polarization ^18^. Upon activation by IL-4/IL-13, STAT6 is phosphorylated and then enters the nucleus, leading to the production of M2-related factors; this process is shown to be regulated by SOCS3 ^24^. In addition, several pattern recognition receptors (PRRs), Dectin-1, 2, 3, and TLR4, 9, are expressed in macrophages and are involved in fungal cell recognition ^25,26^. Among them, the fungal cell wall components, such as chitin, are specifically recognized by TLR9, which then activate a MyD88-dependent signaling pathway that lead to activation of NF-κB and downstream production of anti-inflammatory cytokine IL-10, leading to macrophage M2 polarization ^27^. Perruccio et al. observed that *Aspergillus fumigatus* stimulates the production of IL-10 by stimulating TLR9 in plasmacytoid dendritic cells (pDCs) ^27^. The role SOCS3 and TLR9 play in the promotion of macrophage polarization in *T. marneffei* infection has not been defined.

In this study, we showed that *T. marneffei* evades the antifungal activity of human macrophages by inducing them towards M2 polarization, rendering them permissive for fungal proliferation. We showed that SOCS3 is a positive regulator of M1 polarization in human macrophages, and we showed that *T. marneffei* promotes macrophage M2 polarization by directly down regulating SOCS3 and activating TLR9 pathway.

## Results

### The antifungal response of macrophages against *T. marneffei* infection is associated with macrophage polarization status

It is now known that M1 macrophages enhance their antifungal response by the secretion of antifungal cytokines including TNF-α and IL-1 β, while M2 macrophages dampen their antifungal response by the increased expression of CD163 and CD200R and the secretion of IL-10 ^18,28^. In order to investigate human THP1 macrophages polarization status in response to *T. marneffei,* we challenged the PMA-treated macrophages (M0), PMA/LPS/IFN-γ-treated macrophages (M1), and PMA/IL-4-treated macrophages (M2) with *T. marneffei* spores, respectively. As shown in **Figure 1**, M1 macrophages exhibited the strongest phagocytic function, with significantly higher phagocytic index than M0 or M2 macrophages, while M2 macrophages had the weakest phagocytic function (**Figure 1A**). The *T. marneffei* colony forming units (CFUs) in microdilution spot assay showed that M1 macrophages had the strongest killing activity against *T. marneffei*, whereas M2 macrophages had the weakest killing activity (**Figure 1B**). To confirm whether different treatments of macrophages induced the corresponding polarization status, we measured the levels of a prototypical cytokine produced by M1 macrophages, TNF-α, and a cytokine produced by M2 macrophages, IL-10, in M0, M1, and M2 macrophages, respectively. We showed that M1 macrophages had the highest level of TNF-α and the lowest level of IL-10, while M2 macrophages had the lowest level of TNF-α and the highest level of IL-10 (**Figure 1C**). These findings are consistent with the characteristics of polarized status of M1 macrophages or M2 macrophages.

**FIGURE 1.**
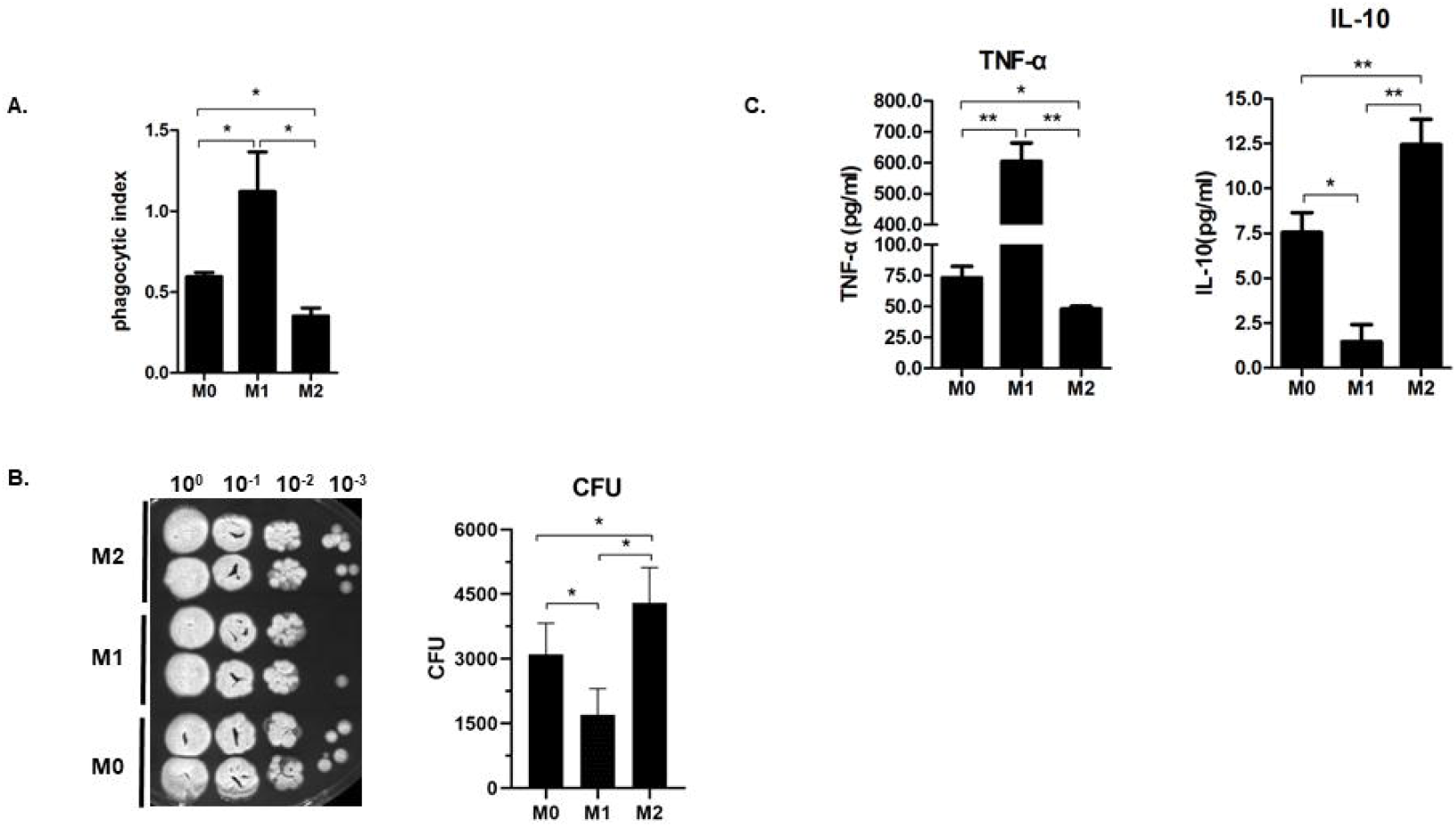
Association between human THP-1 macrophage phenotypes and antifungal response to *T. marneffei*. PMA-treated (M0), PMA/LPS/IFN-γ-treated (M1) or PMA/IL-4-treated (M2) THP-1 macrophages were incubated with *T.marneffei* spores (MOI = 10) for 24h, respectively. (A) Phagocytic index or the number *of T. marneffei* ingested per phagocyte of M0, M1, M2 macrophages. The number of intracellular *T.marneffei* spores were determined by microscope counting. For each group, 100 macrophages were counted and the PI index was calculated as follow: PI = (percentage of phagocytic cells containing ≥ 1 *T. marneffei* spores) × (mean number of *T. marneffei* spores/ phagocytic cells containing *T. marneffei* spores). (B) *T. marneffei* colony forming units (CFU) by microdilution spot assay to assess antifungal activity of M0, M1, M2 macrophages. The culture supernatants and cell lysates were collected, harvested *T.marneffei* by centrifugation. CFUs microdilution spot assay was conducted to measured antifungal ability of M0, M1 or M2 macrophages. (C) Levels of TNF-α and IL-10 in M0, M1 or M2 macrophages as measured by cytometric bead array system (CBA). All data were shown as mean ± SD of the results of at least three independent experiments (*,*p* < 0.05, **,*p* < 0.01, by Student’s t-test).

### *T. marneffei* infection induces human THP-1 macrophages towards M2 polarization

To investigate the impact of *T. marneffei* infection on macrophage polarization, THP-1 macrophages (M0) were infected with *T. marneffei* spores, and the levels of TNF-α and IL-10 were determined by qPCR or cytometric bead array system (CBA). **Figure 2A** showed that *T. marneffei* infection acted in a time-dependent manner to promote the production of IL-10. Although TNF-α was upregulated at 12 hr post-infection, it was significantly down regulated at 48 hr post-infection (**Figure 2A**). We also investigated whether the expression of M2 macrophage markers, CD163 and CD200R, was affected by *T. marneffei* infection of THP-1 macrophages. Flow cytometry results showed that both CD163 and CD200R were significantly upregulated at 12hr, 24hr, and 48hr post-infection (**Figure 2B, C**).

**FIGURE 2.**
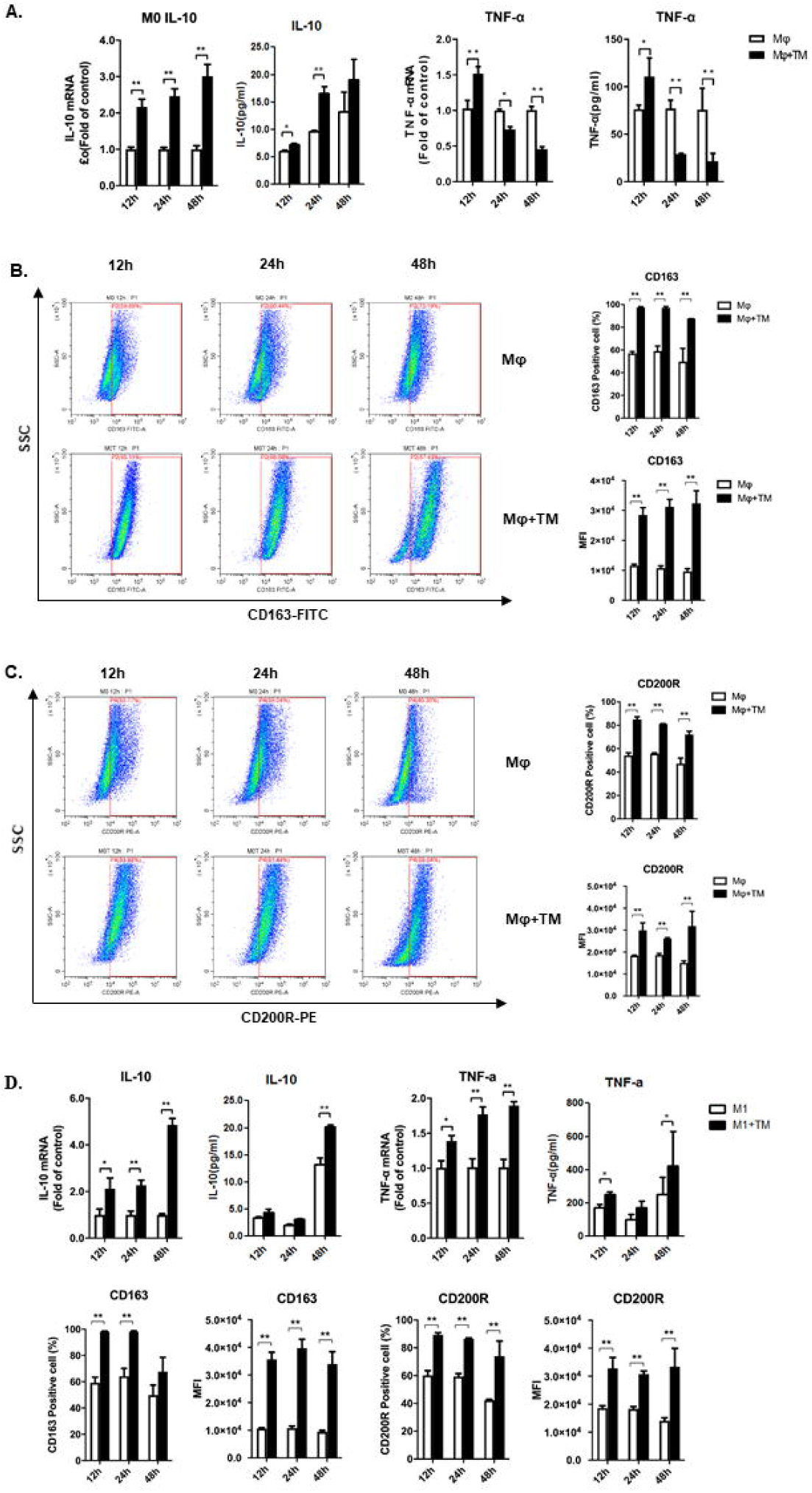
*T. marneffei* infection promotes M2 polarization of human THP-1 macrophage. The human THP-1 macrophages (PMA-treated) or M1 macrophages (PMA/LPS/IFN-γ-treated) were infected with *T.marneffei* spores (MOI = 10) for 12hr, 24hr, or 48hr. (A) TNF-α and IL-10 production in *T. marneffei*-infected THP-1 macrophages at 12hr, 24hr, 48hr post-infection, as measured by qPCR and cytometric bead array system (CBA). (B, C) Levels of CD163 (B) and CD200R (C) in *T. marneffei-infected* macrophages at 12hr, 24hr, 48hr post-infection as measured by flow cytometry. (D) Levels of TNF-α and IL-10 (measured by qPCR and CBA) and the expression of CD163 and CD200R (measured by flow cytometry) in *T. marneffei*-infected M1 macrophages at 12hr, 24hr, 48hr post-infection. All the levels of mRNA expression were normalized to GAPDH and compared with respective timepoint controls. All data were shown as mean ± SD of the results of at least three independent experiments (*,*p* < 0.05, **,*p* < 0.01, by Student’s t-test).

Next, we determined whether *T. marneffei* infection reverses the polarization of M1 macrophages by challenging M1 macrophages with *T. marneffei* spores. We showed that *T. marneffei* infection significantly upregulated the level of IL-10 at both mRNA and protein levels at 12hr, 24hr, and 48hr post-infection (**Figure 2D**). Although the mRNA level of TNF-α was also upregulated by *T. marneffei* infection, the expression of TNF-α at the protein level was not significantly changed by *T. marneffei* infection (**Figure 2D**). These data demonstrated that *T. marneffei* infection induces M2 conversion in human THP-1 M1 macrophages.

### *T. marneffei* infection induces human peripheral blood monocytes/macrophages towards M2 polarization

We tested whether *T. marneffei* infection also affects polarization of human peripheral blood monocytes/macrophages. Human peripheral blood monocytes (PBMCs) isolated from healthy subjects were co-cultured with *T. marneffei* spores, and the expression of CD163 and CD200R in peripheral blood monocytes/macrophages (marked by CD14) were measured by flow cytometry. We found that *T. marneffei* infection induced CD163 and CD200R expression in human PBMCs/macrophages at 12hr and 24hr post-infection (**Figure 3A, B**), suggesting that *T. marneffei* infection also induces M2 polarization of human PBMCs/macrophages.

**FIGURE 3.**
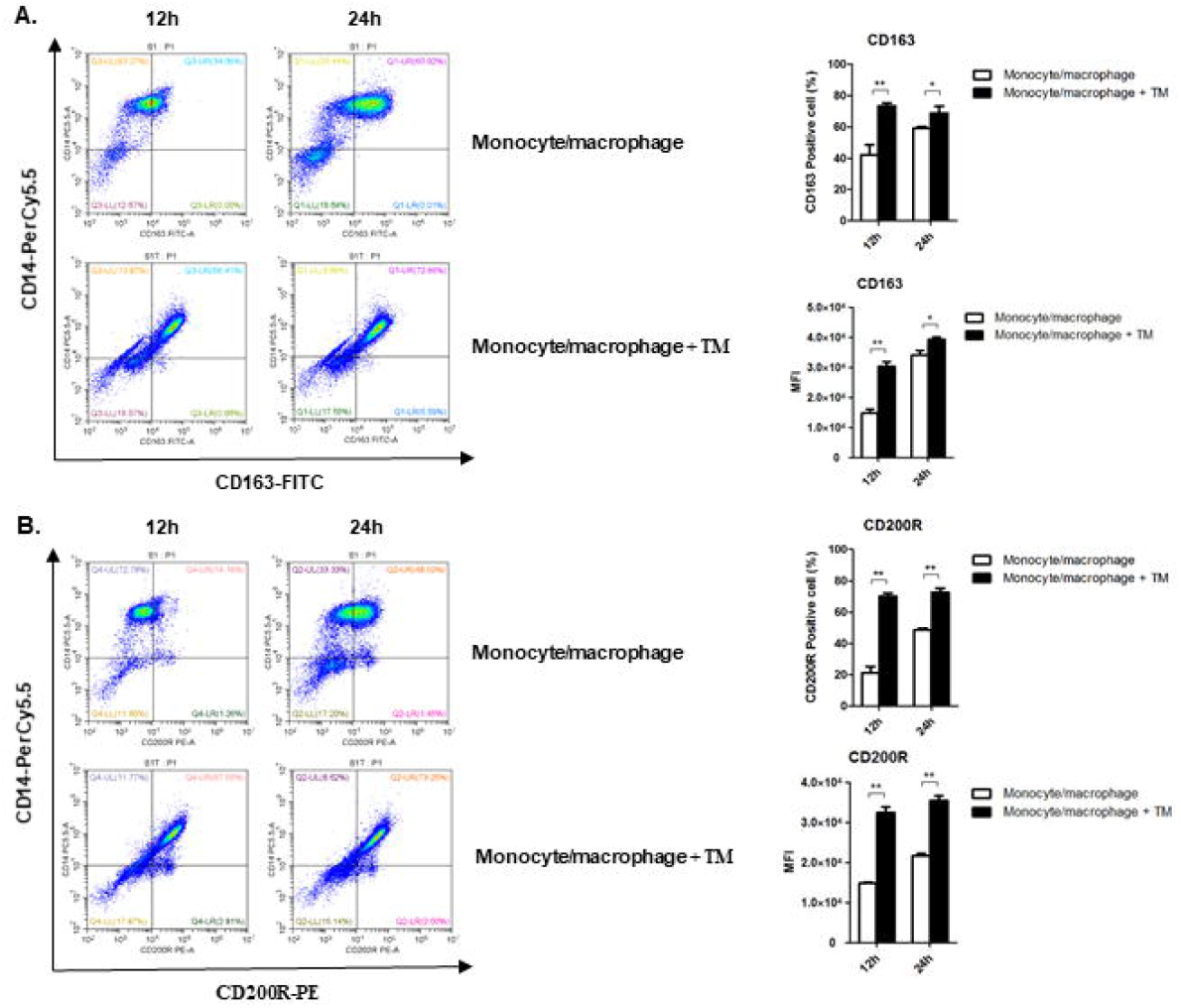
*T. marneffei* infection promotes M2 polarization of human peripheral blood monocytes. Expression of CD163 (A) and CD200R (B) in *T. marneffei-infected* peripheral blood monocytes (PBMC). PBMCs were isolated from peripheral blood of healthy people (n=3), then infected with *T. marneffei* spores at a multiplication of infection (MOI) of 10 for 12hr or 24hr. Human PBMCs were marked by CD14. The expression of CD163 and CD200R was determined by flow cytometry. The data were showed as mean ± SD of results (*,*p* < 0.05, **,*p* < 0.01, by Student’s t-test).

### *T. marneffei* affects SOCS3-STAT6 pathway in THP-1 macrophages

It is well known that SOCS3-STAT6 pathway plays an important role in controlling macrophage polarization. In order to investigate the effect of *T. marneffei* infection on the SOCS3-STAT6 pathway, we infected THP-1 macrophages (M0) with *T. marneffei* spores and measured the levels of SOCS3 mRNA and protein. We found that *T. marneffei* infection upregulated SOCS3 mRNA expression at 12hr post-infection; however, a down-regulated effect was observed at 24hr post-infection (**Figure 4A**). At the protein level, SOCS3 was significantly decreased by *T. marneffei* infection at 12hr, 24hr, 48hr post-infection (**Figure 4B**). Although there is no obvious time-dependent effect at different time points, *T. marneffei-infected* cells expressed significantly lower level of SOCS3 compared with control cells at different time points (**Figure 4B**). Since SOCS3 has inhibitory impact on the phosphorylation of STAT6, we also measured the levels of p-STAT6 in *T. marneffei*-infected cells and control cells. Consistent with the inhibitory effect of *T. marneffei* infection on SOCS3 expression, *T. marneffei* infection led to significantly increased levels of p-STAT6 (**Figure 4C**). Therefore, we showed that the SOCS3-STAT6 pathway was significantly affected during *T. marneffei* infection of human THP-1 macrophages.

**FIGURE 4.**
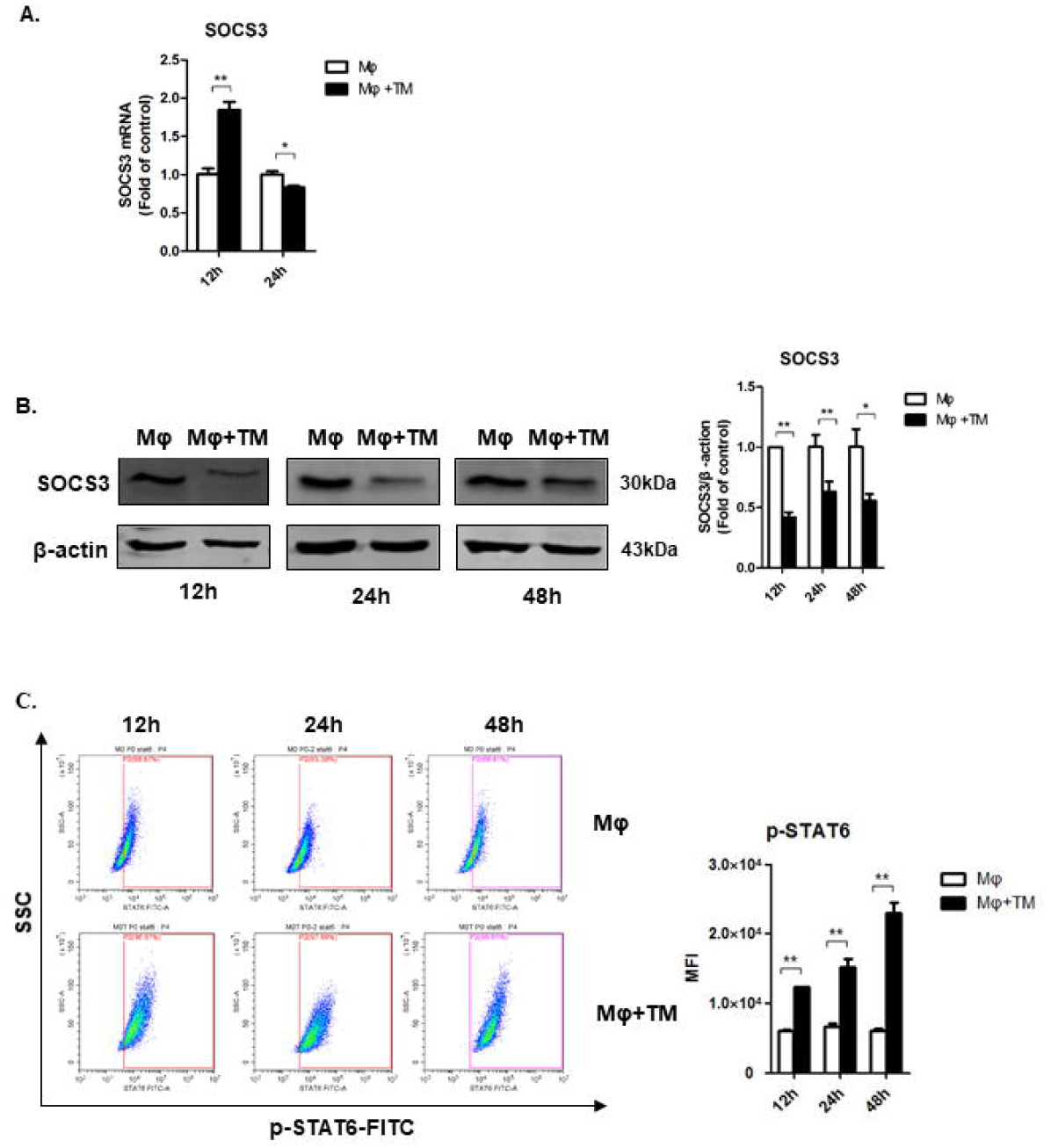
*T. marneffei* infection upregulates SOCS3 and downregulates STAT6 in human THP-1 macrophages. The human THP-1 macrophages (PMA-treated) were infected with *T.marneffei* spores (MOI = 10) for 24hr. (A) Levels of SOCS3 mRNA in *T. marneffei*-infected THP-1 macrophages at 12hr and 24hr post-infection as measured by qPCR. Levels of mRNA expression were normalized to GAPDH and compared with respective timepoint controls. (B) Levels of SOCS3 protein in *T. marneffei*-infected THP-1 macrophages at 12hr, 24hr, 48hr post-infection as measured by Western blot. SOCS3 expression was normalized to ACTIN and compared with respective timepoint controls. (C) Levels of p-STAT6 in *T. marneffei-infected* macrophages at 12hr, 24hr, 48hr post-infection as measured by flow cytometry. All data were showed as mean ± SD of results of at least three independent experiments (*,*p* < 0.05, **,*p* < 0.01, by Student’s t-test).

### SOCS3 plays a critical role in the polarization in THP-1 macrophages

Previous studies have demonstrated the role of SOCS3 in inducing macrophage M1 polarization in rat or mouse models ^17–19^. However, the role of SOCS3 in the polarization of human macrophages has not been demonstrated. To define the role of SOCS3 in human macrophage polarization, a SOCS3-overexpressed THP-1 macrophage cell line was constructed using lentivirus vector, which had twice the level of expression of SOCS3 protein compared to the control cells (**Figure 5A**). The levels of TNF-α and IL-10 in these cells were measured. We showed that SOCS3 overexpression led to significant upregulation of TNF-α at both mRNA and protein level; whereas there was a downregulation of IL-10 expression by SOCS3 overexpression (**Figure 5B**). Furthermore, SOCS3 overexpression inhibited the expression of CD163 and CD200R, two markers of M2 macrophages (**Figure 5C, D**). These results indicate that SOCS3 induces M1 polarization but inhibits M2 polarization in THP-1 macrophages.

**FIGURE 5.**
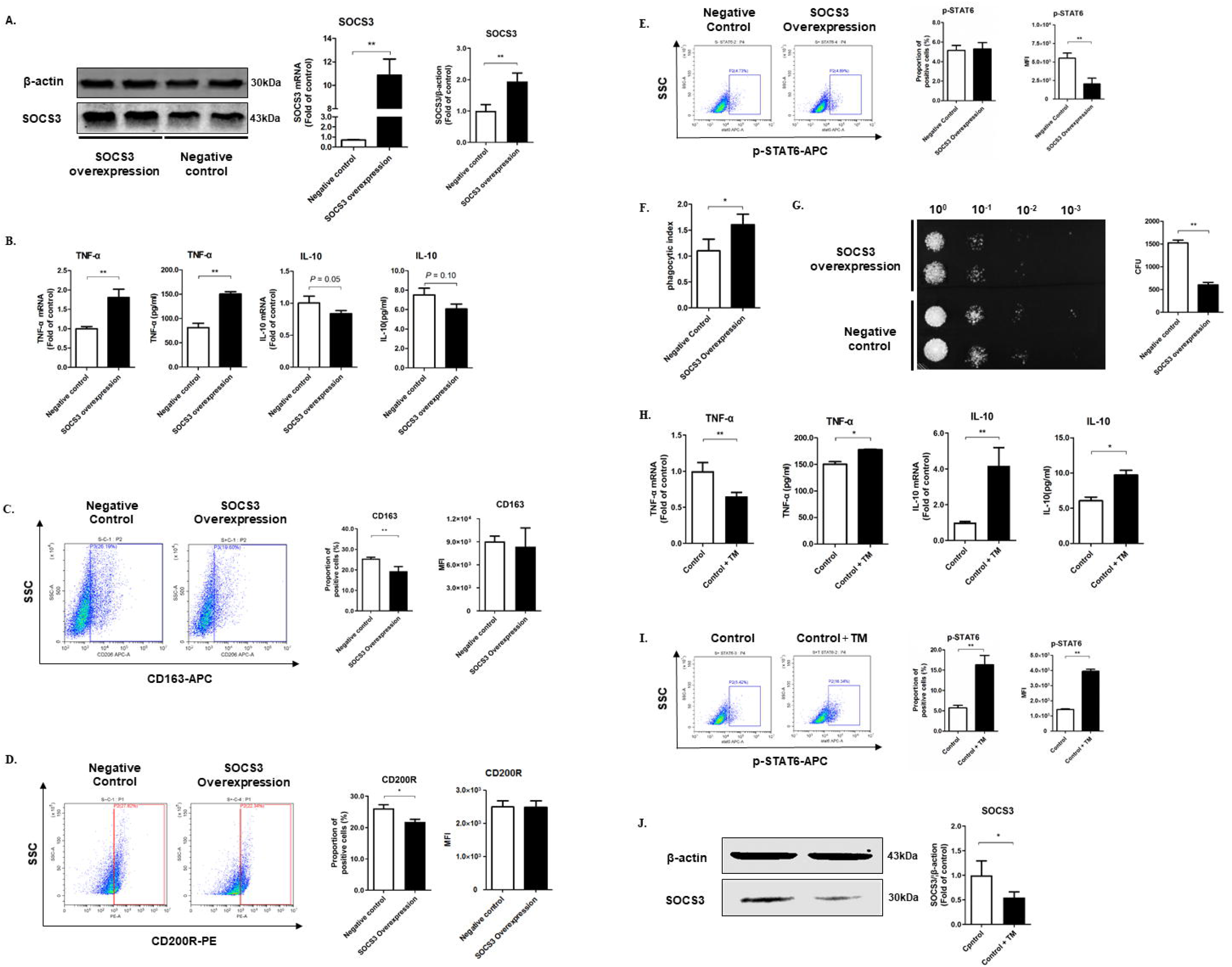
SOCS3 plays an important role in M1 polarization of human THP-1 macrophages in *T. marneffei* infection. (A) Construction of a SOCS3-overexpressed THP-1 macrophage cell line using lentivirus vector with confirmation by qPCR and Western Blot. (B) Levels of TNF-α and IL-10 in SOCS3 - overexpressed THP-1 macrophages and control cells as measured by qPCR and CBA. (C-E) Expression of CD163 (C), CD200R (D) and p-STAT6 (E) in SOCS3-overexpressed THP-1 macrophages and control cells as measured by flow cytometry. (F, G) The effect of SOCS3 overexpression on antifungal function of THP-1 macrophages. SOCS3 overexpression THP-1 macrophages and control cells were incubated with *T.marneffei* spores (MOI =10) for 24h, respectively. (F) Phagocytic activity, as measured by phagocytic index, of SOCS3-overexpressed macrophages. (G) Antifungal activity of SOCS3-overexpressed macrophages as measured by CFUs using microdilution spot assay. (H-J) Effects of *T. marneffei* infection on the expression of TNF-α, IL-10, p-STAT6, SOCS3 in SOCS3-overexpressed THP-1 macrophages. SOCS3-overexpressed THP-1 macrophages were infected with *T. marneffei* spores for 24hr. The levels of TNF-α (H) and IL-10 (H) were measured by qPCR and CBA. The level of p-STAT6 (I) was measured by flow cytometry. The level of SOCS3 (J) was measured by western blot. Levels of mRNA or protein expression were normalized to GAPDH or ACTIN, respectively, and compared with controls.

Next, we measured the effect of SOCS3 on phagocytic capacity and killing activity of macrophages. Compared with control cells, SOCS3-overexpressed THP-1 macrophages had stronger phagocytic activity (**Figure 5E**) and killing activity against *T. marneffei* (**Figure 5F**). By flow cytometry, we observed that STAT6 phosphorylation was significantly inhibited in SOCS3-overexpressed macrophages (**Figure 5G**).

In SOCS3-overexpressed macrophages, *T. marneffei* infection led to significant upregulation of IL-10 compared with control macrophages (**Figure 5H**). Importantly, we showed that *T. marneffei* infection significantly reduced the level of SOCS3 protein and upregulated the level of p-STAT6 in SOCS3-overexpressed macrophages (**Figure 5I, J**). Taken together, we showed that SOCS3 plays an important role in the induction of M1 polarization and in the inhibition of M2 polarization in human THP-1 macrophages, but *T. marneffei* infection partially reversed these effects by the downregulation of SOCS3 expression.

### *T. marneffei* infection induces SOCS3 protein degradation via tyrosine phosphorylation of SOCS3 protein

While the effect of *T. marneffei* on SOCS3 mRNA expression was variable (increased at beginning, then decreased) at different time points (**Figure 4A**), the levels of SOCS3 protein in *T. marneffei-infected* macrophages were consistently lower than control cells at all three time points. This suggests that the regulation of SOCS3 expression by *T. marneffei* infection mainly occurs at the protein level, specifically, via protein degradation. Previous studies have shown that phosphorylation of tyrosine (Tyr204 and/or Tyr221) in the SOCS BOX region of SOCS3 promotes proteasome-mediated degradation of SOCS3 protein ^29^ (**Figure 6A**). We assumed that *T. marneffei* infection causes SOCS3 protein degradation by inducing tyrosine phosphorylation of SOCS3 protein. To test this hypothesis, we used immunoprecipitation and western blot to measure the level of tyrosine phosphorylation of SOCS3 after infecting THP-1 macrophages with *T. marneffei* spores. We showed that SOCS3 tyrosine phosphorylation was significantly increased at 24hr post-infection, accompanied by significantly decreased SOCS3 expression in the infected cells (**Figure 6B**). This indicates that tyrosine phosphorylation of SOCS3 may be an important cause of *T. marneffei*-induced degradation of SOCS3 protein.

**FIGURE 6.**
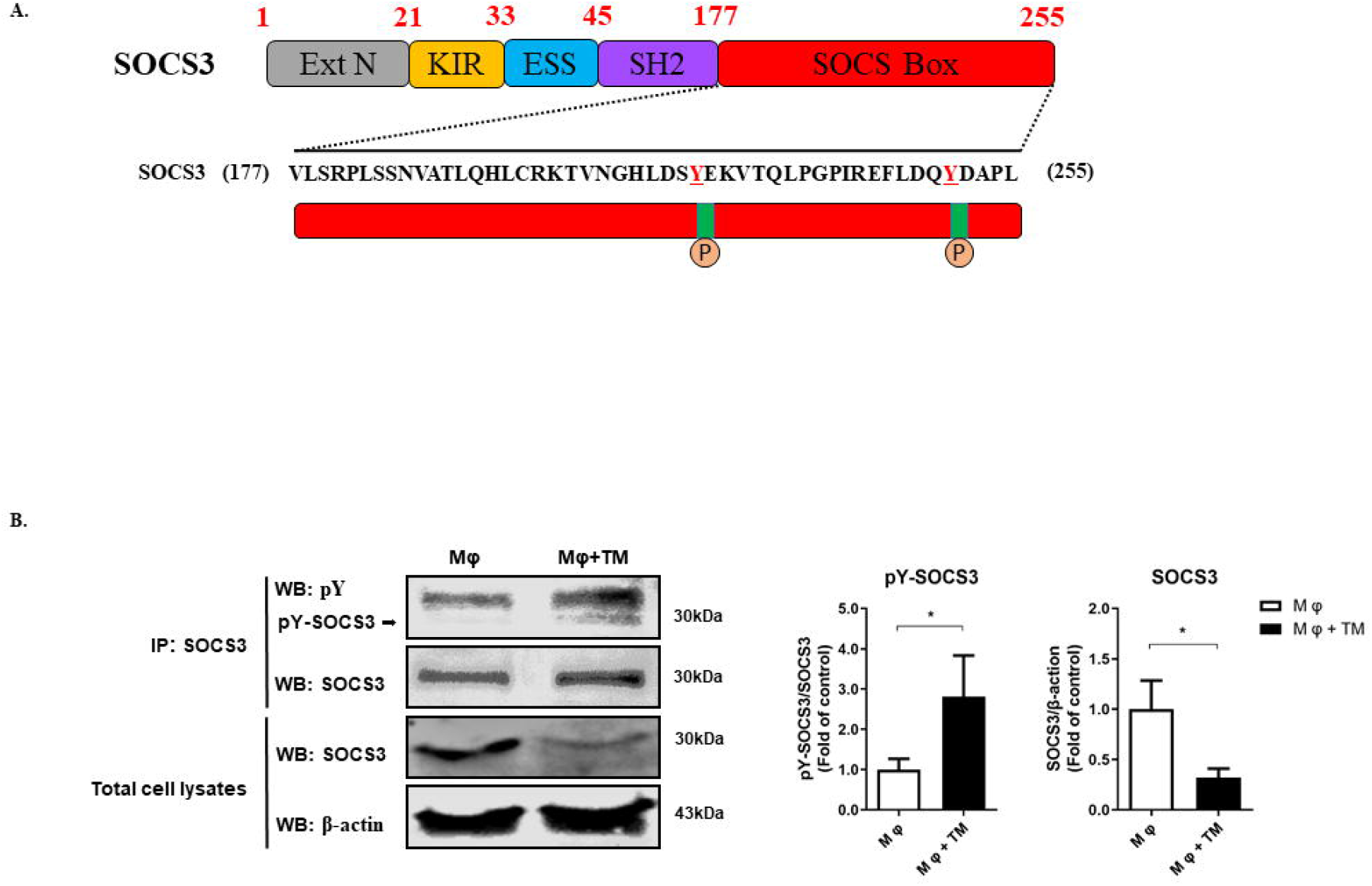
*T. marneffei* infection induces tyrosine phosphorylation of SOCS3 protein thereby enhancing SOCS3 protein degradation in THP-1 macrophages. The human THP-1 macrophages (PMA-treated) were infected with *T.marneffei* spores (MOI = 10) for 24hr. (A) The structure of SOCS3 protein and the amino acid sequence of SOCS BOX region. Tyrosine (Tyr204 and/or Tyr221) in the SOCS BOX region can be phosphorylated to initiate proteasome-mediated degradation of SOCS3 protein. (B) The effect of *T.marneffei* infection on tyrosine phosphorylation of SOCS3. The macrophages were lysised and were co-incubated with capture SOCS3 antibody for immunoprecipitation. Western blot was used to detect the levels of SOCS3 tyrosine phosphorylation and SOCS3 in *T.marneffei-infected* or uninfected macrophages. The relative pY-SOCS3/SOCS3 ratios were calculated and shown as fold of control. And the relative SOCS3/β-actin ratios were calculated and shown as fold of control (without *T.marneffei* infection). All data were showed as mean ± SD of results of three independent experiments (*, *p* < 0.05, **, *p* < 0.01, by Student’s t-test).

### TLR9 pathway is involved in *T. marneffei*-induced M2 polarization of THP-1 macrophages

Previous studies have shown that SOCS3 plays an important role in the regulation of M1 polarization ^19^. However, *T. marneffei* reduces SOCS3 expression does not mean that SOCS3 degradation is sufficient for the induction of M2 polarization. Other factors may contribute to *T. marneffei*-induced M2 polarization. TLR9 pathway has been reported to be involved in M2 polarization of macrophages, and TLR9 is a key receptor that recognizes *T. marneffei* infection. We hypothesize that TLR9 is involved in *T. marneffei*-induced macrophage M2 polarization. To test this hypothesis, we suppressed TLR9 receptor of THP-1 macrophages via its antagonist, ODN-4084F, and then infected the cells with *T. marneffei* and measured the production of TNF-α and IL-10. We showed that the suppression of TLR9 pathway completely blocked IL-10 upregulation in response to *T. marneffei* infection. However, it insignificantly increased the expression of TNF-α (**Figure 7A**). The suppression of TLR9 pathway partially inhibited *T. marneffei*-induced upregulation of CD163 and CD200R (**Figure 7B, C**). Importantly, the role of TLR9 pathway in *T. marneffei-induced* M2 polarization was evidenced by the fact that TLR9 antagonist improved the killing activity of THP-1 macrophages against *T. marneffei* (**Figure 7D**). These data showed that TLR9 mediates M2 polarization of THP-1 macrophages in response to *T. marneffei* infection.

**FIGURE 7.**
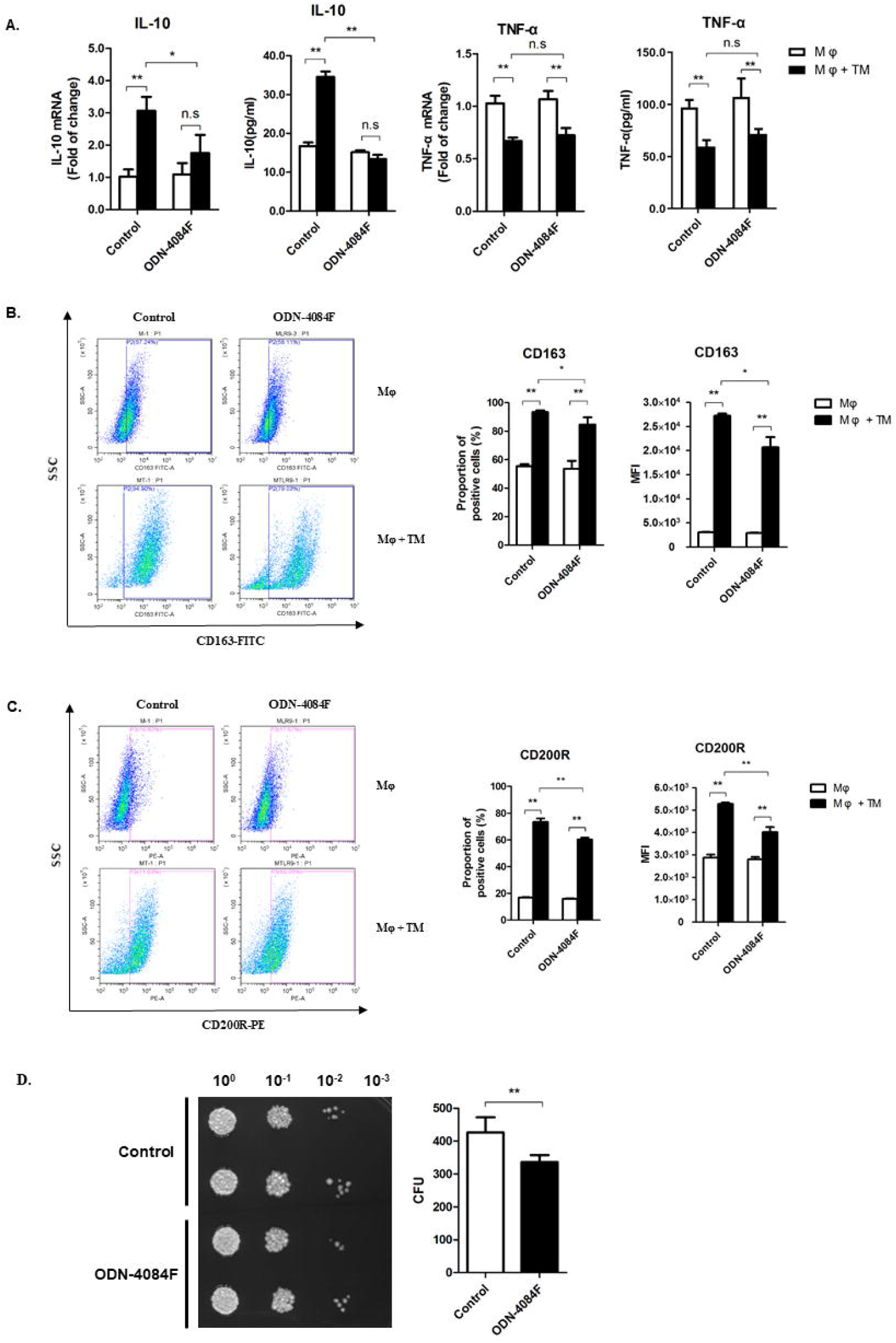
*T. marneffei* infection mediates M2 polarization of THP-1 macrophages by TLR9 pathway. THP-1 macrophages were treated with 1 μ M of ODN-4084F or control ODN for 24hr, then infected or uninfected with *T. marneffei* spores (MOI = 10) for 24hr. (A) Levels of TNF-α and IL-10 as measured by qPCR and CBA. All the levels of mRNA expression were normalized to GAPDH and compared with respective timepoint controls. (B, C) Expression of CD163 (B) and CD200R (C) as measured by flow cytometry. (D) The effect of TLR9 on antifungal function of macrophages. Culture supernatants and cell lysates were collected, harvest *T.marneffei* by centrifugation. And the CFUs microdilution spot assay were conducted to detect the killing ability of macrophages to *T.marneffei.* All data were showed as mean ± SD of results of three independent experiments (*,*p* < 0.05, **, *p* < 0.01, n.s, no significant difference, by Student’s t-test).

We further explored the relationship between SOCS3 and TLR9 in macrophage M2 polarization. First, we suppressed TLR9 pathway in SOCS3-overexpressed THP-1 macrophages, then infected the cells with *T. marneffei* spores and measured the production of TNF-α and IL-10. Suppression of TLR9 pathway or/and SOCS3 overexpression significantly reduced the production of IL-10 in response to *T. marneffei* infection, and partially blocked the upregulation of TNF-α (**Figure 8A**). In addition, TLR9 suppression or/and SOCS3 overexpression partially inhibited *T. marneffei*-mediated upregulation of CD163 and CD200R (**Figure 8B, C**). Similarly, both TLR9 suppression and SOCS3 overexpression enhanced the killing activity of macrophages against *T. marneffei*, respectively (**Figure 8D**). Moreover, TLR9 suppression in SOCS3-overexpressed macrophages showed the strongest ability to kill *T. marneffei* (**Figure 8D**). Collectively, we demonstrated that *T. marneffei-induced* M2 polarization of human THP-1 macrophages is additively regulated by SOCS3 and TLR9 pathway.

**FIGURE 8.**
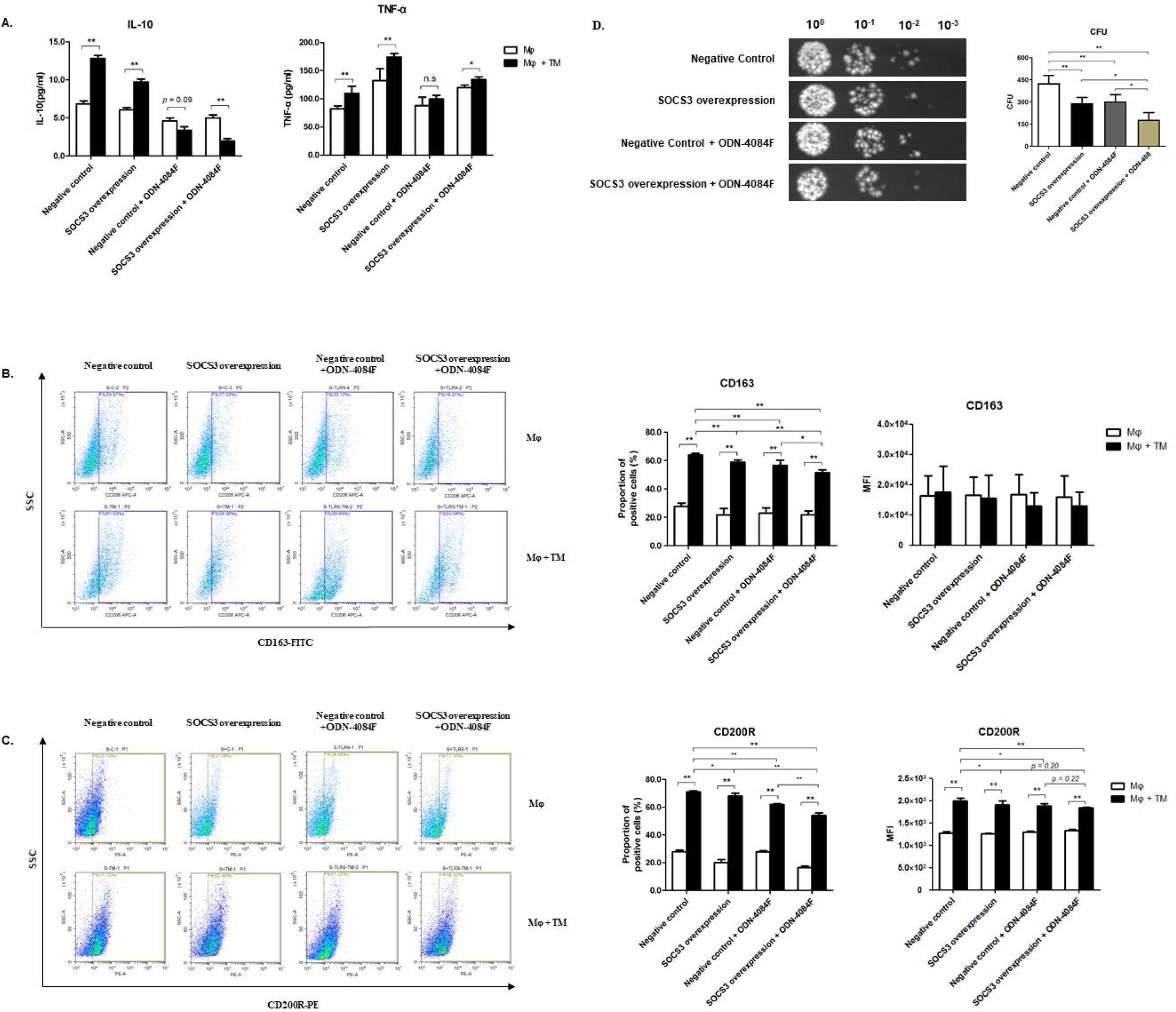
TLR9 and SOCS3 additively mediate M2 polarization in *T. marneffei-induced* THP-1 macrophages. SOCS3-overexpressed macrophages or control cells were treated with 1 μ M of ODN-4084F or control ODN for 24hr, then infected or uninfected with *T. marneffei* spores (MOI = 10) for 24hr. (A) Levels of TNF-α and IL-10 were detected by CBA. (B, C) Expression of CD163 (B) and CD200R (C) are measured by flow cytometry. (D) The effect of TLR9 on antifungal function of SOCS3 overexpression macrophages and control cells. Culture supernatants and cell lysates were collected, harvest *T.marneffei* by centrifugation. And the CFUs microdilution spot assay were conducted to detect the killing ability of macrophages to *T.marneffei.* All data were showed as mean ± SD of results of three independent experiments (*,*p* < 0.05, **,*p* < 0.01, n.s, no significant difference, by Student’s t-test).

## Discussion

In this study, we demonstrated that *T. marneffei* resisted the direct fungicidal effects of macrophage by inducing macrophage M2 polarization. We showed that SOCS3-STAT6 and TLR9 pathways directly participated in this process. Specifically, *T. marneffei* infection downregulated the level of SOCS3 protein, thereby disinhibiting the effect of SOCS3 on STAT6 phosphorylation and facilitating macrophage M2 polarization. We observed that SOCS3 protein is actually a positive regulator of the M1 polarization in human macrophages. Moreover, M2 polarization of macrophages induced by *T. marneffei* infection is also mediated by TLR9 activation.

Macrophage polarization has emerged as a key role in antifungal activity of macrophages. Similar to other fungal and intracellular pathogens, we observed the ability of *T. marneffei* to shift human macrophage polarization status from M1 to M2 to evade the antifungal activity of THP-1 macrophages. The intracellular environment of macrophages is harsh by design, including oxidative stress, heat stress and nutrient deprivation ^30,31^. *T. marneffei* has been shown to develop strategies to withstand these stresses. These include challenging oxidative stress via nuclear localization of transcription factors and signal transduction via phosphorylation, responding to heat stress by initiating yeast morphogenesis, and using the glyoxylate cycle to overcome the dilemma of nutritional deprivation ^32,33^. Inducing M2 polarization of macrophage is another effective strategy for *T. marneffei* to resist antifungal killing of macrophages, rendering them permissible to fungal persistence and dissemination. Of note, *T. marneffei*-induced M2 polarization was not only observed in human macrophage cell line (THP-1), but also in human PBMCs/macrophages (**Figure 2**), partially implying that *T. marneffei*-induced macrophage M2 polarization may be a common phenomenon in human macrophages. In addition, our results suggested that there appears to be a population of CD163-low macrophages arising in *T. marneffei-induced* group at 48h (**Figure 2B**), which was similar to the results of other studies^34,35^. We speculate that macrophages may have a negative feedback regulation mechanism, and there may be an upper limit to the degree of M2 polarization of macrophages.

Our finding that SOCS3 shifts macrophage polarization from M2 to M1 phenotype is similar to a previous study ^17^. Using adoptive transfer of SOCS3-silenced macrophages in a mouse peritonitis model, the investigators demonstrated that SOCS3 drives the production of M1-induced cytokines (TNF-α, IL-1β), while reducing the expression of M2-induced IL-10 ^19^. Our finding is also consistent with another study in rat bone marrow derived macrophages, in which silencing of SOCS3 led to down-regulation of the M1 polarization gene ^23^. Interestingly, there are species-specific and cells-specific differences in the role of SOCS3 in driving macrophage polarization ^18^. Specifically, SOCS3 has been shown to be a positive regulator of M1 polarization in mouse bone-marrow-derived macrophages and in bone-marrow-derived macrophages from glioma patients, but is a positive regulator of M2 polarization in human THP-1 macrophages and rat bone marrow derived macrophages ^20–22,36^. Overall, a unifying role of SOCS3 is macrophage polarization across species has not been established, and further research are need to better define the role and mechanism of SOCS3 in the regulation of macrophage polarization against fungal infections.

As an important member of SOCS protein family, SOCS3 is a cytokine-inducible negative regulator of cytokine signaling pathways. Since the IL4/IL13-driven JAK-STAT6 pathway directly promotes macrophage M2 polarization ^17,18^, we hypothesize that the JAK-STAT6 pathway is involved in *T. marneffei-induced* M2 polarization of macrophages. Our results confirmed this hypothesis showing that *T. marneffei* infection activated JAK-STAT6 pathway by downregulating the level of SOCS3. Our results further found that tyrosine phosphorylation of SOCS3 could be selectively activated by *T. marneffei* infection, and tyrosine phosphorylation within the SOCS3 box region directly induced SOCS3 degradation ^29,37,38^. Our data therefore provided a complete chain of evidence that *T. marneffei* infection degrades SOCS3 protein, thereby disinhibits the effect of SOCS3 on STAT6 phosphorylation, and facilitates M2 polarization of macrophages which is beneficial for the survival of *T. marneffei* in THP-1 macrophages.

Besides the JAK-STAT6 pathway, TLR9 has been shown to be involved in M2 polarization ^27^. TLR9 receptor mediates cellular response to cytidine phosphate guanosine (CpG) in fungal DNA to trigger an innate immune response ^39–41^. TLR9 directly recognizes chitin of fungal cell wall, thereby increases the production of IL-10 ^16^. In the present study, the inhibition of TLR9 pathway by ODN-4084F significantly blocked the upregulation of IL-10 and enhanced phagocytosis and killing activity of human THP-1 macrophages upon *T. marneffei* infection. This suggests that chitin on *T. marneffei* may be involved in the blocking of the TLR9 pathway. The down regulation of CD163 and CD200R by ODN-4084F further support our insertion that TLR9 pathway is important for the regulation of M2 polarization in THP-1 macrophage.

There are several limitations in the present study. First, only one cell line, THP-1 macrophage, was used. Considering the species-specific and cell-specific characteristics of SOCS3, it is unclear whether this mechanism is reproducible in other macrophage cell lines. Second, the SOCS3 knockdown cell line needs to be further studied the role of SOCS3 in macrophage polarization. Third, our in vitro experiments do not simulate the immune microenvironment in vivo. In vivo infection, the body’s immune cells, such as CD4^+^ T cells, neutrophils, NK cells, and macrophages interact with each other to coordinate an antifungal response. Therefore, the question whether *T. marneffei* can evade immune killing by inducing macrophage M2 polarization requires further verification in in vivo experiments.

Collectively, our data lead us to propose that *T. marneffei* avoids macrophage killing by modulating the SOCS3-STAT6 and TLR9 pathways to induce M2 polarization in human THP-1 macrophages. Our study reveals a novel mechanism by which *T. marneffei* evades the innate immune response, which may provide a therapeutic target for inhibition of infection and dissemination in *T. marneffei* infection.

## Materials and Methods

### Cell line and Peripheral blood mononuclear cells (PBMCs)

The human monocytic cell line THP-1 was purchased from Chinese Academy of Sciences Cell Bank. The THP-1 monocytes were stimulated with 50 ng/ml PMA for 72 hours for monocytes differentiation to macrophages, as previously described ^42^. A SOCS3-overexpressed THP-1 cell line was generated using lentivirus construction strategy. The lentivirus vector containing green fluorescent protein (GFP)-SOCS3 sequences was constructed by Genechem Company (Shanghai, China). Peripheral blood mononuclear cells (PBMCs) were isolated from healthy volunteers. Briefly, blood sample was collected from healthy subjects (n=3). PBMCs were isolated by Ficoll-Paque Plus (GE Healthcare) density centrifugation according to the manufacturer’s instructions. The study, including the recruitment of healthy volunteers, was approved by the Institutional Review Board and Human Ethics Committee of Guangxi Medical University, and the informed consents were obtained from all the subjects. THP-1 cells and PBMCs were cultured in RPMI 1640 medium containing 10% FBS, 100 U/mL penicillin, and 100 μg/mL streptomycin. THP-1 differentiated macrophage was maintained in 10% heat-inactivated FBS in DMEM with 100 U/mL penicillin and 100 μg/mL streptomycin. The cells were cultured at 37 °C and 5% CO_2_. The medium was changed every 3 days.

### Fungal strain and media

Experiments were conducted using *T. marneffei* strain L0, which was separated from an HIV and *T. marneffei* co-infected patient. *T. marneffei* was identified by both standard culture morphology and by PCR-based sequence analysis of the 16S-23S rRNA internal transcribed spacer region (ITS). *T. marneffei* was plated onto *Potato dextrose agar* (PDA) agar at 27°C. The conidia were harvested by suspension in PBS after incubation for 7 to 10 days on PDA. The purified conidia were separated by filtering through sterile glass wool. The conidial suspension with the concentration of 10^7^ conidia/ml was prepared for subsequent experiments.

### *T. marneffei* infection

The purified conidia suspended in RPMI 1640 or DMEM medium with 10% FBS were added to PBMCs or THP-1 macrophages, respectively (MOI = 10), as previously described ^43^.

### Phagocytosis assay

Phagocytosis assay was carried out as previously described ^14^. In brief, *T. marneffei* spores were stained with calcofluor white (CFW; 10μg/ml; Sigma-Aldrich) for 10 min at room temperature, and macrophages were stained with CFSE (100μM; Dojindo) for 30 min. Then the CFSE-positive macrophages were infected with CFW-positive *T. marneffei* spores for 24hr. Fluorescence was observed with Invitrogen EVOS FL Auto Cell Imaging System. Phagocytic index (PI) was calculated as follow: PI = (percentage of phagocytic cells containing ≥ 1 *T. marneffei* spores) × (mean number of *T. marneffei* spores/ phagocytic cells containing *T. marneffei* spores).

### *T. marneffei* survival assay

In brief, THP-1 macrophages were incubated with T. *marneffei* (MOI = 10) for 24h at 37°C and 5% CO_2_. For ODN-4084-F related experiments, cells were seeded on 96-well plates at 1 ×10^5^ cells per well, rest of which were seeded on 24-well plates at 3 × 10^5^ cells per well, ensure that cells were seeded at 90% confluence into different plates^44,45^. The supernatant was collected, and the cells were lysed in sterile water to release the fungus. Four gradient serial dilutions (10^0^, 10^1^, 10^2^, and 10^3^) were performed and plated onto YPD agar. Numbers of fungal colony-forming units (CFUs) were counted after 24 h incubation at 30°C.

### Quantitative Real-Time PCR (qPCR)

qPCR was carried out as previously described ^46^. In brief, total cellular RNA was extracted using Universal RNA Extraction Kits (Takara). Then the Reverse Transcription Kit (Takara) was used to reversely transcribe RNA to cDNA, according to the manufacturer’s instructions. Gene expression was examined with a StepOne Plus real-time PCR system (Life Technologies) with SYBR Green PCR Master Mix (Takara). The values were normalized to GAPDH expression.

### Western blot

Western blot was carried out as previously described ^46^. Briefly, total cell lysates were prepared using radioimmune precipitation assay (RIPA) buffer (MultiSciences Biotech) with 1% protease inhibitor cocktail (Asvio Technology). The primary antibodies used in this study were listed as follows: rabbit-anti-β-actin (1:1000) (CST), rabbit-anti-SOCS3 (1:1000) (CST), mouse-anti-Phosphotyrosine (1:500) (Millipore). The secondary antibodies were horseradish peroxidase-conjugated goat-anti-rabbit IgG (1:15000) (LI-COR Biosciences) or goat-anti-mouse IgG (1:15000) (LI-COR Biosciences), respectively. The immunoreactive bands were visualized by Odyssey CLx Infrared Imaging System (LI-COR Biosciences). The densitometric analysis of blots was performed by Image Studio Ver 5.2 software (LI-COR Biosciences). The values were normalized to those of control β-actin.

### Flow Cytometry

Flow Cytometry was carried out according to the manufacturer’s instructions (BD Biosciences). In brief, for cell surface staining, the cells were incubated with Human BD Fc Block™ at room temperature for 10min, then antibodies were added, and incubation maintained for 30 min in the absence of light at 4°C. For intracellular staining, the cells were fixed with incubating pre-warmed BD Cytofix^TM^ Fixation Biffer for 10 min at 37°C. Then the cells were incubated with BD Phosflow™ Perm/Wash Buffer III for 30 min at room temperature. At last, antibodies were added, and incubation maintained for 60 min at 4°C protected from light. For Cytometric Bead Assay (CBA), the cells were incubated with beads at room temperature for 1hr, then antibodies were added, and incubation maintained for 2 hr at room temperature protected from light. Fluorescence was detected with Beckman CytoFLEX FCM, and the data was analyzed by Beckman CytExpert 2.0 Software. The antibodies used in this study were listed as follows: BD Pharmingen™ FITC Mouse Anti-Human CD163, BD Pharmingen™ Alexa Fluor^®^ 647 Mouse Anti-Human CD163, eBioscience™ PE Mouse Anti-Human CD200R, and BD Phosflow™ Alexa Fluor^®^ 488 Mouse Anti-Stat6 (pY641). The CBA Flex Sets used in this study were BD™ Rat TNF Flex Set and BD™ Mouse IL-10 Flex Set.

### Immunoprecipitation

Immunoprecipitation was conducted using PureProteome^TM^ Protein A/G Mix Magnetic Beads (Millipore). Briefly, the macrophages were lysed using lysis buffer (20 mM Tris-HCl pH 7.5, 150mM NaCl, 1mM Na2EDTA, 1% Triton, 1mM EGTA, 1mM NA3VO4) supplemented with 1% protease/phosphataes inhibitor cocktail (CST), and incubated on ice for 5 minutes. The supernatant was collected by centrifugation at 14,000g for 10 minutes, and co-incubated with capture SOCS3 antibody (5μg/10^7^ cells; Abcam) at 4°C overnight. The antibody-antigen complex was then added to the beads and incubated for 30 min at room temperature. Finally, adding the appropriate elution buffer for denaturing elution after washing the complex. The samples were heated to 95-100 °C for 5 minutes, and then cooled on ice. Immunoprecipitated proteins were analyzed by western blot with a rabbit-anti-SOCS3 antibody (Abcam). To eliminate the interference of rabbit lgG heavy and light chains, Mouse Anti-rabbit lgG (Conformation Specific) (CST), which only reacts with native lgG and doesn’t bind to the denatured and reduced rabbit lgG heavy and light chains, were used as the secondary antibody for western blot.

### Ethical statement

All blood donors gave informed written consent for use of their blood samples for scientific purposes.

### Statistical analysis

All experimental groups were performed in triplicates. Experimental groups were compared using Student’s *t* test for pair-wise comparisons and using one-way analysis variance (ANOVA) for multiple group comparisons. Data were shown as mean ± standard deviation (SD).

## DATA AVAILABILITY STATEMENT

The datasets generated for this study are available on request to the corresponding author.

## ETHICS STATEMENT

The studies involving human participants were reviewed and approved by Ethics and Human Subjects Committee of Guangxi Medical University (Ethical Review No. 20200096). The patients/participants provided their written informed consent to participate in this study.

## AUTHOR CONTRIBUTIONS

WW, CN, JH performed most of the experiments and analyzed the data. GW, JL, JH, OZ, NZ, and BL also provided experimental data or generated proviral constructs. BL, NZ, JJ, TL, HL, and LY provided reagents and expertise. WW, CN, JH, TL, HL, and JJ conceived and designed the experiments. WW, CN, and JH wrote the initial draft manuscript. All authors have seen, edited, and approved the final version of the manuscript.

## ACKNOWLEDGMENTS

We would like to express our gratitude to all participants involved in this study.

## REFERENCES

1. Limper A H, Adenis A, Le THarrison T S. Fungal infections in HIV/AIDS. The Lancet. Infectious diseases 2017; 17: e334–e343. 10.1016/S1473-3099(17)30303-1

2. Nongnuch V, Cooper C R, Fisher M CThira S. Penicillium marneffei infection and recent advances in the epidemiology and molecular biology aspects. Clinical Microbiology Reviews 2006; 19: 95–110.

3. Le T, Kinh N V, Cuc N T K, Tung N L N, Lam N T, Thuy P T T, Cuong D D, Phuc P T H, Vinh V H, Hanh D T H, Tam V V, Thanh N T, Thuy T P, Hang N T, Long H B, Nhan H T, Wertheim H F L, Merson L, Shikuma C, Day J N, Chau N V V, Farrar J, Thwaites G, Wolbers MInvestigators I. A Trial of Itraconazole or Amphotericin B for HIV-Associated Talaromycosis. N Engl J Med 2017; 376: 2329–2340. 10.1056/NEJMoa1613306

4. Jiang J, Meng S, Huang S, Ruan Y, Lu X, Li J Z, Wu N, Huang J, Xie Z, Liang B, Deng J, Zhou B, Chen X, Ning C, Liao Y, Wei W, Lai J, Ye L, Wu FLiang H. Effects of Talaromyces marneffei infection on mortality of HIV/AIDS patients in southern China: a retrospective cohort study. Clin Microbiol Infect 2019; 25: 233–241. 10.1016/j.cmi.2018.04.018

5. Chan J F, Lau S K, Yuen K YWoo P C. Talaromyces (Penicillium) marneffei infection in non-HIV-infected patients. Emerg Microbes Infect 2016; 5: e19. PMC4820671; 10.1038/emi.2016.18

6. Thuy L, Wolbers M, Nguyen H C, Vo M Q, Nguyen T C, Nguyen P H L, Pham S L, Kozal M J, Shikuma C M, Day J NFarrar J. Epidemiology, Seasonality, and Predictors of Outcome of AIDS-Associated Penicillium marneffei Infection in Ho Chi Minh City, Viet Nam. Clinical Infectious Diseases 2011; 52: 945–952. 10.1093/cid/cir028

7. Hu Y, Zhang J, Li X, Yang Y, Zhang Y, Ma JXi L. Penicillium marneffeiInfection: An Emerging Disease in Mainland China. Mycopathologia 2013; 175: 57–67.

8. Kawila R, Chaiwarith RSupparatpinyo K. Clinical and laboratory characteristics of penicilliosis marneffei among patients with and without HIV infection in Northern Thailand: a retrospective study. BMC infectious diseases 2013; 13: 1–5. Artn 464.1186/1471-2334-13-464

9. Altmeier SLeibundgut-Landmann S. Immunity to Fungal Infections, (2017).

10. Murray P J. Macrophage Polarization. Annu Rev Physiol 2017; 79: 541–566. 10.1146/annurev-physiol-022516-034339

11. Dai X, Mao C, Lan X, Chen H, Li M, Bai J, Deng J, Liang Q, Zhang J, Zhong X, Liang Y, Fan J, Luo HHe Z. Acute Penicillium marneffei infection stimulates host M1/M2a macrophages polarization in BALB/C mice. BMC Microbiol 2017; 17: 177. PMC5563047; 10.1186/s12866-017-1086-3

12. Chen R, Xi L, Huang X, Ma T, Ren HJi G. Effect of Jun N-terminal kinase 1 and 2 on the replication of Penicillium marneffei in human macrophages. Microb Pathog 2015; 82: 1–6. 10.1016/j.micpath.2015.03.014

13. Ellett F, Pazhakh V, Pase L, Benard E L, Weerasinghe H, Azabdaftari D, Alasmari S, Andrianopoulos ALieschke G J. Macrophages protect Talaromyces marneffei conidia from myeloperoxidase-dependent neutrophil fungicidal activity during infection establishment in vivo. PLoS Pathog 2018; 14: e1007063. PMC6010348; 10.1371/journal.ppat.1007063

14. Wagener J, MacCallum D M, Brown G DGow N A. Candida albicans Chitin Increases Arginase-1 Activity in Human Macrophages, with an Impact on Macrophage Antimicrobial Functions. mBio 2017; 8: e01820–01816. PMC5263244; 10.1128/mBio.01820-16

15. Feng X, Kang Y, Hang Z, Piao Z, Yin H, Ran D, Xia JShi L. Akt1 comprised antibacterial response through regulating macrophage polarization. Journal of Infectious Diseases 2013.

16. Price J VVance R E. The macrophage paradox. Immunity 2014; 41: 685–693. 10.1016/j.immuni.2014.10.015

17. Wilson H M. SOCS Proteins in Macrophage Polarization and Function. Front Immunol 2014; 5: 357. PMC4112788; 10.3389/fimmu.2014.00357

18. McCormick S MHeller N M. Regulation of Macrophage, Dendritic Cell, and Microglial Phenotype and Function by the SOCS Proteins. Front Immunol 2015; 6: 549. PMC4621458; 10.3389/fimmu.2015.00549

19. Arnold C E, Whyte C S, Peter G, Barker R N, Rees A JWilson H M. A critical role for suppressor of cytokine signalling 3 in promoting M1 macrophage activation and function in vitro and in vivo. Immunology 2013; 141: 96–110.

20. Hongwei Q, Holdbrooks A T, Yudong L, Reynolds S L, Yanagisawa L LBenveniste E N. SOCS3 deficiency promotes M1 macrophage polarization and inflammation. Journal of Immunology 2012; 189: 3439–3448.

21. Chunguang Y, Ward P A, Ximo WHongwei G. Myeloid depletion of SOCS3 enhances LPS-induced acute lung injury through CCAAT/enhancer binding protein δ pathway. Faseb Journal Official Publication of the Federation of American Societies for Experimental Biology 2013; 27: 2967–2976.

22. Yasukawa H, Ohishi M, Mori H, Murakami M, Chinen T, Aki D, Hanada T, Takeda K, Akira S, Hoshijima M, Hirano T, Chien K RYoshimura A. IL-6 induces an anti-inflammatory response in the absence of SOCS3 in macrophages. Nat Immunol 2003; 4: 551–556. 10.1038/ni938

23. Yu L, Stewart K N, Eileen B, Marek C J, Kluth D C, Rees A JWilson H M. Unique expression of suppressor of cytokine signaling 3 is essential for classical macrophage activation in rodents in vitro and in vivo. Journal of Immunology 2008; 180: 6270–6278.

24. Jiang M, Zhang W W, Liu P, Yu W, Liu TYu J. Dysregulation of SOCS-Mediated Negative Feedback of Cytokine Signaling in Carcinogenesis and Its Significance in Cancer Treatment. Front Immunol 2017; 8: 70. PMC5296614; 10.3389/fimmu.2017.00070

25. Maldonado SFitzgerald-Bocarsly P. Antifungal Activity of Plasmacytoid Dendritic Cells and the Impact of Chronic HIV Infection. Front Immunol 2017; 8: 1705. PMC5723005; 10.3389/fimmu.2017.01705

26. Saijo SIwakura Y. Dectin-1 and Dectin-2 in innate immunity against fungi. Int Immunol 2011; 23: 467–472. 10.1093/intimm/dxr046

27. Jeanette W, Malireddi R K S, Lenardon M D, Martin K B, Simon V, Maccallum D M, Tilo B, Martin S, Netea M GThirumala-Devi K. Fungal chitin dampens inflammation through IL-10 induction mediated by NOD2 and TLR9 activation. Plos Pathogens 2014; 10: e1004050.

28. Murray P J, Allen J E, Biswas S K, Fisher E A, Gilroy D W, Goerdt S, Gordon S, Hamilton J A, Ivashkiv L B, Lawrence T, Locati M, Mantovani A, Martinez F O, Mege J L, Mosser D M, Natoli G, Saeij J P, Schultze J L, Shirey K A, Sica A, Suttles J, Udalova I, van Ginderachter J A, Vogel S NWynn T A. Macrophage activation and polarization: nomenclature and experimental guidelines. Immunity 2014; 41: 14–20. PMC4123412; 10.1016/j.immuni.2014.06.008

29. Williams J J, Munro K MPalmer T M. Role of Ubiquitylation in Controlling Suppressor of Cytokine Signalling 3 (SOCS3) Function and Expression. Cells 2014; 3: 546–562. PMC4092859; 10.3390/cells3020546

30. Pongpom M, Vanittanakom P, Nimmanee P, Jr C R CVanittanakom N. Adaptation to macrophage killing byTalaromyces marneffei. Future Science Oa 2017; 3: FSO215.

31. Brown A J, Haynes KQuinn J. Nitrosative and oxidative stress responses in fungal pathogenicity. Curr Opin Microbiol 2009; 12: 384–391. PMC2728829; 10.1016/j.mib.2009.06.007

32. Moye-Rowley W S. Regulation of the transcriptional response to oxidative stress in fungi: similarities and differences. Eukaryot Cell 2003; 2: 381–389. PMC161443; 10.1128/ec.2.3.381-389.2003

33. Tiwari S, Thakur RShankar J. Role of Heat-Shock Proteins in Cellular Function and in the Biology of Fungi. Biotechnol Res Int 2015; 2015: 132635. PMC4736001; 10.1155/2015/132635

34. Zeng M Y, Pham D, Bagaitkar J, Liu J Y, Otero K, Shan M, Wynn T A, Brombacher F, Brutkiewicz R R, Kaplan M HDinauer M C. An efferocytosis-induced, IL-4-dependent macrophage-iNKT cell circuit suppresses sterile inflammation and is defective in murine CGD. Blood 2013; 121: 3473–3483. 10.1182/blood-2012-10-461913

35. Jung S B, Choi M J, Ryu D, Yi H S, Lee S E, Chang J Y, Chung H K, Kim Y K, Kang S G, Lee J H, Kim K S, Kim H J, Kim C S, Lee C H, Williams R W, Kim H, Lee H K, Auwerx JShong M. Reduced oxidative capacity in macrophages results in systemic insulin resistance. Nat Commun 2018; 9: 1551. PMC5908799; 10.1038/s41467-018-03998-z

36. McFarland B C, Marks M P, Rowse A L, Fehling S C, Gerigk M, Qin HBenveniste E N. Loss of SOCS3 in myeloid cells prolongs survival in a syngeneic model of glioma. Oncotarget 2016; 7: 20621–20635. PMC4991480; 10.18632/oncotarget.7992

37. Serge H, Paul F, Ulrike S, Meena H, Mcvicar D W, Heinrich P C, Johnston J ACacalano N A. Tyrosine phosphorylation disrupts elongin interaction and accelerates SOCS3 degradation. Journal of Biological Chemistry 2003; 278: 31972–31979.

38. Sommer U, Schmid C, Sobota R M, Lehmann U, Stevenson N J, Johnston J A, Schaper F, Heinrich P CHaan S. Mechanisms of SOCS3 phosphorylation upon interleukin-6 stimulation - Contributions of Src- and receptor-tyrosine kinases. Journal of Biological Chemistry 2005; 280: 31478–31488. 10.1074/jbc.M506008200

39. Ramirez-Ortiz Z G, Specht C A, Wang J P, Lee C K, Bartholomeu D C, Gazzinelli R TLevitz S M. Toll-like receptor 9-dependent immune activation by unmethylated CpG motifs in Aspergillus fumigatus DNA. Infect Immun 2008; 76: 2123–2129. PMC2346696; 10.1128/IAI.00047-08

40. Botos I, Segal D MDavies D R. The structural biology of Toll-like receptors. Structure 2011; 19: 447–459. PMC3075535; 10.1016/j.str.2011.02.004

41. Nakamura K, Miyazato A, Xiao G, Hatta M, Inden K, Aoyagi T, Shiratori K, Takeda K, Akira S, Saijo S, Iwakura Y, Adachi Y, Ohno N, Suzuki K, Fujita J, Kaku MKawakami K. Deoxynucleic acids from Cryptococcus neoformans activate myeloid dendritic cells via a TLR9-dependent pathway. Journal of immunology (Baltimore, Md.: 1950) 2008; 180: 4067–4074. 10.4049/jimmunol.180.6.4067

42. Komoda H, Shiraki A, Oyama J, Nishikido TNode K. Azelnidipine Inhibits the Differentiation and Activation of THP-1 Macrophages through the L-Type Calcium Channel. Journal of Atherosclerosis and Thrombosis 2018; 25: 690–697. 10.5551/jat.41798

43. Gomez P, Hackett T L, Moore M M, Knight D ATebbutt S J. Functional genomics of human bronchial epithelial cells directly interacting with conidia of Aspergillus fumigatus. BMC Genomics 2010; 11: 358. PMC2897809; 10.1186/1471-2164-11-358

44. Mo J, Xie Q, Wei WZhao J. Revealing the immune perturbation of black phosphorus nanomaterials to macrophages by understanding the protein corona. Nat Commun 2018; 9: 2480. PMC6018659; 10.1038/s41467-018-04873-7

45. Danelishvili L, Everman JBermudez L E. Mycobacterium tuberculosis PPE68 and Rv2626c genes contribute to the host cell necrosis and bacterial escape from macrophages. Virulence 2016; 7: 23–32. PMC4871676; 10.1080/21505594.2015.1102832

46. Meihua R, Hui C, Bingyu L, Weibo L, Junjun J, Jiegang H, Chuanyi N, Ning Z, Bo ZYanyan L. Alcohol-induced autophagy via upregulation of PIASy promotes HCV replication in human hepatoma cells. Cell Death & Disease 2018; 9: 898-.

